# Comparison of Mortality and Viral Load in Rainbow Trout (*Oncorhynchus mykiss*) infected with Infectious Pancreatic Necrosis Virus (IPNV) Genogroups 1 and 5

**DOI:** 10.1101/719104

**Authors:** David Tapia, Agustín Barría, José M. Yáñez

## Abstract

Infectious Pancreatic Necrosis Virus (IPNV) is the etiological agent of a highly contagious disease that affects farmed salmonids. IPNV isolates have been phylogenetically classified into eight genogroups, of which two are present in Chile, genogroups 1 and 5. Here we compare the mortality rate caused by isolates from both genogroups in rainbow trout (*Oncorhynchus mykiss*) fry to determine if there is an association between host susceptibility and phylogenetic characterization of IPNV. Fish were challenged by immersion with one of four isolates (two of each genogroup) and mortality curves were assessed after 30 days. Viral load was measured in all mortalities and in live fish sampled at 1, 7 and 20 days post-infection. Although mortality was low throughout the challenge, differences were found between fish infected with different isolates. Both isolates from genogroup 1, caused greater cumulative mortalities than either of the isolates from Genogroup 5. When combined, the overall mortality rate of fish challenged with genogroup 1 isolates was significantly higher than those infected with genogroup 5. However, viral load was lower on trout infected with genogroup 1 isolates. These results suggest that rainbow trout are more susceptible to IPNV isolates from genogroup 1 than genogroup 5.

## INTRODUCTION

Infectious Pancreatic Necrosis (IPN) is a highly contagious viral disease that causes great economic losses to trout and salmon aquaculture worldwide. The etiological agent of the disease, IPN virus (IPNV), a non-enveloped, double-stranded RNA (dsRNA) virus which affects the main salmonid species cultured worldwide, i.e. Atlantic salmon (*Salmo salar*) and rainbow trout (*Oncorhynchus mykiss*). The disease is transmitted horizontally as well as vertically, and after an outbreak surviving fish can become asymptomatic carriers of the virus, representing a risk to their offspring and other susceptible fish (Bootland et al., 1991). Salmonids are more susceptible to IPNV during their first feeding fry stage in fresh water, where prevalence of the disease is high, and as post-smolts shortly after transfer to the sea (Jarp et al., 1995; Evensen & Santi, 2008). During an IPN outbreak, mortality rates can vary greatly from insignificant to almost 100%, and these differences have been ascribed to several environmental, viral and host-related factors. Host genetic variability associated to IPNV mortality has been confirmed in Atlantic salmon with the identification of a major-effect quantitative trait loci (QTL), associated with mortality levels in fresh water and seawater (Houston *et al*., 2008; Moen et al., 2009), and the subsequent identification of epithelial cadherin associated with resistance to IPNV (Moen et al. 2015). In rainbow trout, resistance against IPNV has also shown significant genetic variation (Flores-Mara et al., 2017; Yoshida et al., 2019). Furthermore, a recent study detected a marker which moderately explains the genetic variance for this trait (Rodríguez et al., 2019).

Amino acids 217 and 221 of the IPNV capsid protein, VP2, are associated with differences in virulence and replication rates between isolates (Santi et al., 2004; Song et al. 2005). Thus, IPNV isolates with amino acids Thr_217_ and Ala_221_ are considered highly virulent and Pro_217_ and Thr_221_ as almost avirulent (Song et al., 2005). However, these markers have only been studied in IPNV type Spjarup (Sp, genogroup 5), and do not always correlate with field mortality rates seen in IPN outbreaks (Ruane et al., 2015; Tapia et al., 2015).

IPNV belongs to the genus *Aquabirnavirus*, from the *Birnaviridae* family. Its genome consists of two dsRNA segments: segment A, which encodes a viral capsid protein (VP2), an internal protein (VP3), a viral protease (VP4), and a non-structural protein (VP5); and segment B, which encodes the RNA polymerase (VP1) (Dobos, 1995). *Aquabirnaviruses* were initially classified based on antiserum neutralization assays directed against three IPNV reference serotypes: VR299, Sp and Ab (Macdonald & Gower, 1981). Subsequently, a standard serological classification scheme was proposed, and the strains were divided into serogroups A and B. The former serogroup contains the majority of IPNV isolates associated with the disease and are grouped into 9 serotypes, A1-A9 (Hill & Way, 1995). More recently, *Aquabirnaviruses* were classified according to their nucleotides and deduced amino acid sequences. Phylogenetic analysis of the coding region of the VP2 protein from 28 aquatic birnavirus isolates revealed that the nine reference strains of serogroup A were grouped into six genogroups, some of which were composed of several genotypes (Blake et al., 2001). These genogroups are highly correlated with the serological classification and geographic origin of the strains, for example, genogroup 5 corresponds to serogroup A2 and includes several isolates from Europe such as Sp, Fr10 and N1, while genogroup 1 correlates to serotype A1 and includes strains from the United States, such as West Buxton (WB), VR-299 and Buhl (Blake et al., 2001). Following this approach, a seventh genogroup comprised only by Japanese aquabirnaviruses was proposed by Nishizawa et al. (2005). An Aquabirnavirus isolated from *Oncorhynchus mykiss* in Australia was classified in an eighth genogroup (McCowan et al., 2015; Mohr et al., 2015).

Chile is currently the largest producer of rainbow trout worldwide (FAO 2019); and it was in this species that IPNV was isolated for the first time in the country (McAllister & Reyes 1984). The isolate corresponded to serotype VR-299 or 1A (Espinoza et al., 1985, McAllister & Reyes, 1984), and it was later classified in genogroup 1 (Eissler et al., 2011). During the late nineties, IPN disease was confirmed in Atlantic salmon farms and since then the virus has spread through the country’s different farming areas. Only two IPNV genogroups have been reported in the country, genogroup 1 (North American origin) and genogroup 5 (European origin), with the latter being more dominant and widely dispersed (Mutoloki & Evensen, 2011, Eissler *et al*., 2011, Calleja *et al*., 2012; Tapia *et al*., 2015; Manríquez *et al*. 2017).

More recently, Torres *et al*. (2016) analysed data from Chilean isolates and suggested a host-specific relationship between the reported genogroups and the salmonid species cultivated in the country, since isolates belonging to genogroup 5 were mainly isolated from *S. salar*, while IPNV genogroup 1 was mostly isolated from *O. mykiss* or *O. Kisutch*. However, to date there are no experimental studies aimed to evaluate a relationship between these two genogroups and the differential susceptibility between host species. Thus, the objective of this study was to compare the mortality rates and viral load in rainbow trout infected with isolates from genogroups 1 and 5 to confirm the likely association between this species and the phylogenetic classification of IPNV.

## MATERIALS AND METHODS

### Virus isolates and fish

Four IPNV isolates obtained from field outbreaks in Chile were used for the experimental challenges in this study. The isolates belong to the strain collection of the Laboratory of Virology of the University of Valparaíso and were molecularly characterized previously by sequencing both segments of the virus genome according to Jorquera *et al*. (2016). The four isolates represent the two IPNV genogroups present in the country: two isolates from genogroup 1 (type strain WB), herein referred to as WB1 and WB2, respectively; and two from genogroup 5 (type strain Sp), herein referred to as SP1 and SP2, respectively. All isolates were propagated and titrated in the Chinook Salmon Embryo (CHSE-214) cell line. For virus propagation, cells were grown in Leibovitz (L-15) culture medium, supplemented with 10% fetal bovine serum (FBS, HyClone) and 50 μg·mL-1 gentamicin. Then the cells were infected with first passages of the isolates at a multiplicity of infection (MOI) of 0.001 FF/cell and incubated at 15 °C until a massive cytopathic effect (CPE) was observed (around 3 to 5 days post-infection). The cells were then subjected to two cycles of freezing and thawing and centrifuged at 3000 × g for 15 min at 4°C. Supernatants were collected, aliquoted and immediately stored at −80°C until titration or challenge inoculation.

Titration was done twice, between two to three days after the isolates were propagated and prior to the inoculation of the fish. For standard end-point titration, aliquots of the isolates were thawed and serially diluted in L-15 medium and inoculated into confluent monolayers of CHSE-214 cells grown on 96-well plates. The cells were incubated at 15 °C for a week and then observed under an inverted light microscope. The infected wells with clear ECP were counted and the TCID_50_ was calculated according to the Reed & Muench (1938) method.

Rainbow trout fry (∼ 0.9 g) were provided by the Rio Blanco hatchery (Los Andes, Chile) and transferred to the aquaculture facilities of the “Laboratorio Cerrillos”, Veterquimica S.A. (Santiago, Chile). Prior to the challenge test, the fish were sanitarily checked for the presence of IPNV, *Renibacterium Salmoninarum, Piscirickettsia salmonis, Flavobacterium psychrophilum* and *F. columnare* by qPCR. All of these diagnostic analyses were carried out at Veterquimica diagnostic laboratory. Fish were held at 10-15 °C and fed daily on commercial feed until the day preceding virus exposure.

### Experimental challenges and Sampling

In order to compare the mortality caused by the IPNV isolates in rainbow trout, 1,440 fish were challenged during January 2018. Prior to challenge, fish were acclimatized for 22 days and then allocated into eight 0.02 m^3^ aerated challenge tanks, with ∼180 individuals per tank. The experimental design included two tank replicates per treatment. Once distributed in their respective tanks, fish were challenged by immersion using the four IPNV isolates. For this, duplicate groups of fish were exposed to each isolate at 1 x 10^5^ TCID50/mL^-1^. Thereafter, fish were maintained at 10 °C and mortality was recorded daily until the trial ended 30 days post-infection. Dead fish were weighed and the entire viscera, including kidney, was sampled to confirm the presence of IPNV and to estimate viral load, following the procedures recommended by the World Organisation for Animal Health (OIE 2006). Mortality samples were maintained in L-15 medium at 4 °C until homogenization. Additionally, six fish were taken randomly from each tank and sacrificed by benzocaine overdose (BZ-20; Veterquimica, Chile) at days 1, 7 and 20 post-infection. The fish were weighed and tissue samples were taken and stored in RNAlater® solution (Ambion, USA) at −80 °C until use. All procedures for challenges and sampling were approved by the Comite de Bioetica Animal, Facultad de Ciencias Veterinarias y Pecuarias, Universidad de Chile (Certificate N° 17086-VET-UCH).

### IPNV testing and viral load

Samples from mortality and survivor fish were tested to confirm the presence of IPNV and to estimate viral load. A RT-qPCR assay for the VP1 region of the virus (Eissler *et al*. 2011) was used for confirmation of the presence of the virus in both mortality and survivor fish samples, while an indirect fluorescent antibody test (IFAT) targeting the VP2 protein of the virus (Espinoza & Kuznar 2002) was used only on samples from mortalities. For the latter, homogenization was done in 1 mL of L-15 medium using bead beating in the Minlys (Bertin) homogenizer. The homogenates were pelleted at 2000 × g (15 min at 4°C) in a MIKRO 22R centrifuge (Hettich Zentrifugen), and the supernatants were collected and stored at −80°C until use. For the IFAT assay the methodology described by Espinoza & Kuznar (2002) was followed with slight modifications. Briefly, 100 μL of each supernatant was serially diluted in L-15 medium and inoculated into duplicate confluent monolayers of CHSE-214 cells grown on 24-well plastic plates with 12 mm circular glass coverslips. After 24 h, the cell monolayers were fixed with cold methanol for 10 min, rinsed with PBS 1× buffer, and incubated with a polyclonal antibody against VP2 IPNV during 1 h at room temperature. Cells were rinsed again, and a secondary anti-rabbit antibody (Sigma) conjugated with fluorescein isothiocyanate (FITC) was used to label the infected cells (1/100 in PBS). After rinsing the cells for a third time, the circular glass cover-slips containing the cell monolayers were mounted on glass slides using an anti-fade mounting solution (DakoCytomation) and visualized using an epifluorescence microscope (Olympus BX60). For the RT-qPCR assay, total RNA was extracted from 200 μL of each supernatant from the mortality samples with the E.Z.N.A._TM_ Total RNA Kit I (Omega Bio-tek) according to the manufacturer’s instructions. The extracted RNA was eluted with molecular biology grade water and stored at −80°C until use. In the case of samples taken from alive fish at different timepoints (1, 7 and 20 days), tissues (maintained in RNAlater) were homogenized in 1 mL of TRIzol reagent (Invitrogen) by bead beating, and total RNA was extracted following the standard TRIzol RNA isolation procedure.

Concentration and purity of the extracted total RNA was determined by measuring the absorbance ratio at 260 nm over 280 nm using a spectrophotometer (MaestroNano, Maestrogen). To ensure that contamination was strictly controlled during the RNA extraction process, a negative control using molecular biology grade water was included. The extracted RNA was reversely transcribed and amplified by a one-step RT-qPCR using a 48-well plate real-time PCR system Step-One (Applied Biosystems). The sets of primers and probes used for the VP1 and for the Elongation Factor −1 alpha (ELF1α) RT-qPCR assays are shown in Table 1. The AgPath-IDTM One-Step RT-PCR Kit (Applied Biosystems) was used for the amplification of the VP1 protein in segment B. Reaction was carried out in a 15 μL reaction volume containing 7.5 μL of RT-PCR Buffer (2X), 1.35 μL of each forward and reverse primers (0.9 μM), 0.3 μL of the VP1 Taqman probe (0.2 μM), 0.6 μL of RT-PCR Enzyme Mix (25X) and 2 μL of total RNA as template. The thermal profile used was 48°C for 10 min for reverse transcription, pre-denaturation at 95°C for 10 min, followed by 40 cycles of denaturation at 95°C for 15 s and annealing/extension at 59°C for 45 s. In case of ELF1α amplification was carried out in a 15 μL reaction volume containing 7.5 μL of 2X Brilliant III Ultra-Fast SYBR® Green QRT-PCR Master Mix (Stratagene), 0.75 μL of each forward and reverse primers (0.5 μM), 0.8 μL of RT/RNAse block, 0.2 μL of ROX (0.3 μM) as passive reference and 2 μL of total RNA as template.

The thermal profile used was 50°C for 5 min for reverse transcription, pre-denaturation at 95°C for 3 min, followed by 40 cycles of denaturation at 95°C for 5 s and annealing/extension at 60°C for 10 s. Finally, a melting curve analysis from 70°C to 95°C was performed. The detection limit and the efficiency of the assays were evaluated using 10-fold dilutions of total RNA from the virus and from non-infected rainbow trout. The amplification efficiencies were 103% for the IPNV RT-qPCR targeting segment B (Taqman) and 100% for the ELF1 RT-qPCR (SYBR green), and the cut-off Ct values were 30.8 and 32.4, respectively.

**Table 1.**
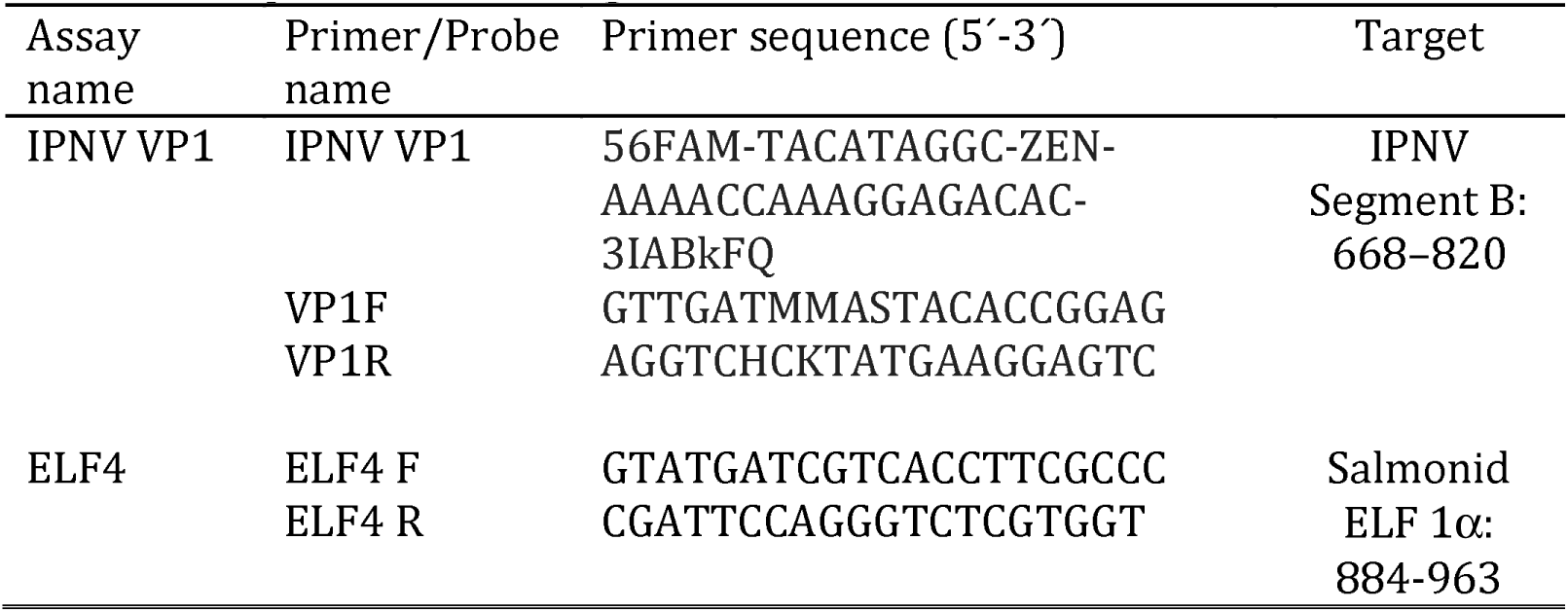
Primers and probes used for the RT-qPCR assays to identify and quantify IPNV in samples from challenged individuals.

### Statistical analysis

Kaplan-Meier mortality curves were calculated using the GraphPad Prism 7 software (GraphPad Software Inc., Ja Lolla, CA, USA). For this analysis, mortalities were considered the result of IPNV infection only in cases when IPNV presence was confirmed by IFAT and/or RT-qPCR, while live fish sampled at different time points were not taken into account. Cumulative mortality rates over time between fish infected with different isolates and genogroups were compared using the Log-rank test (p < 0.05). In order to compare the results and evaluate the agreement between the two diagnostic methods used, the Kappa statistic was calculated. The scale used to interpret the Kappa statistic was as follows: below 0.01 less than chance agreement, 0.01–0.20 slight agreement, 0.21–0.40 fair agreement, 0.41–0.60 moderate agreement, 0.61–0.80 substantial agreement and 0.81–0.99 indicate almost perfect agreement (Viera & Garrett 2005). For the estimation of viral load in dead and survivor fish samples by RT-qPCR, relative expression of IPNV VP1 mRNA was calculated using the Pfaffl method (Pfaffl 2001) which accounts for differences in PCR efficiency. ELF1α was used as a housekeeping gene for normalization, and the sample with the lowest VP1 gene expression (*i.e.* highest Ct value) was set as calibrator. Finally, differences in viral load were analyzed using an analysis of covariance (ANCOVA), in which the dependent variable (viral load) was explained by a lineal model including tank nested to genogroup as factor and time of death and body weight of fish as covariate.

## RESULTS

### Mortality

Mortality due to IPNV started at day 4 post-challenge in fish infected with either WB1 or WB2. In fish challenged with isolates from genogroup 5, mortality began on day 11 (SP2) and day 13 (SP1) post infection. Cumulative mortality at day 30 post infection was low in all groups challenged, but especially in fish infected with isolates from genogroup 5, with final cumulative mortalities of 0.67% and 1% for SP1 and SP2, respectively. Fish infected with genogroup 1 isolates showed a higher cumulative mortality, with 2.33% for WB2 and 7.33% for WB1. Overall, trout infected with isolates belonging to genogroup 5 showed a total cumulative mortality of 0.84%. The total cumulative mortality reached 4.73% for genogroup 1. Kaplan–Meier (Log-Rank) analysis showed that the mortality curves of fish infected with the four isolates were significantly different (*p* <0.0001), and that isolate WB1 had a significantly higher mortality rate over time than either of the two isolates from genogroup 5 (Figure 1). Furthermore, when comparing the two genogroups, the mortality rate of trout infected with genogroup 1 isolates was significantly higher than of those challenged with isolates from genogroup 5 (*p* <0.0001).

**Figure 1.**
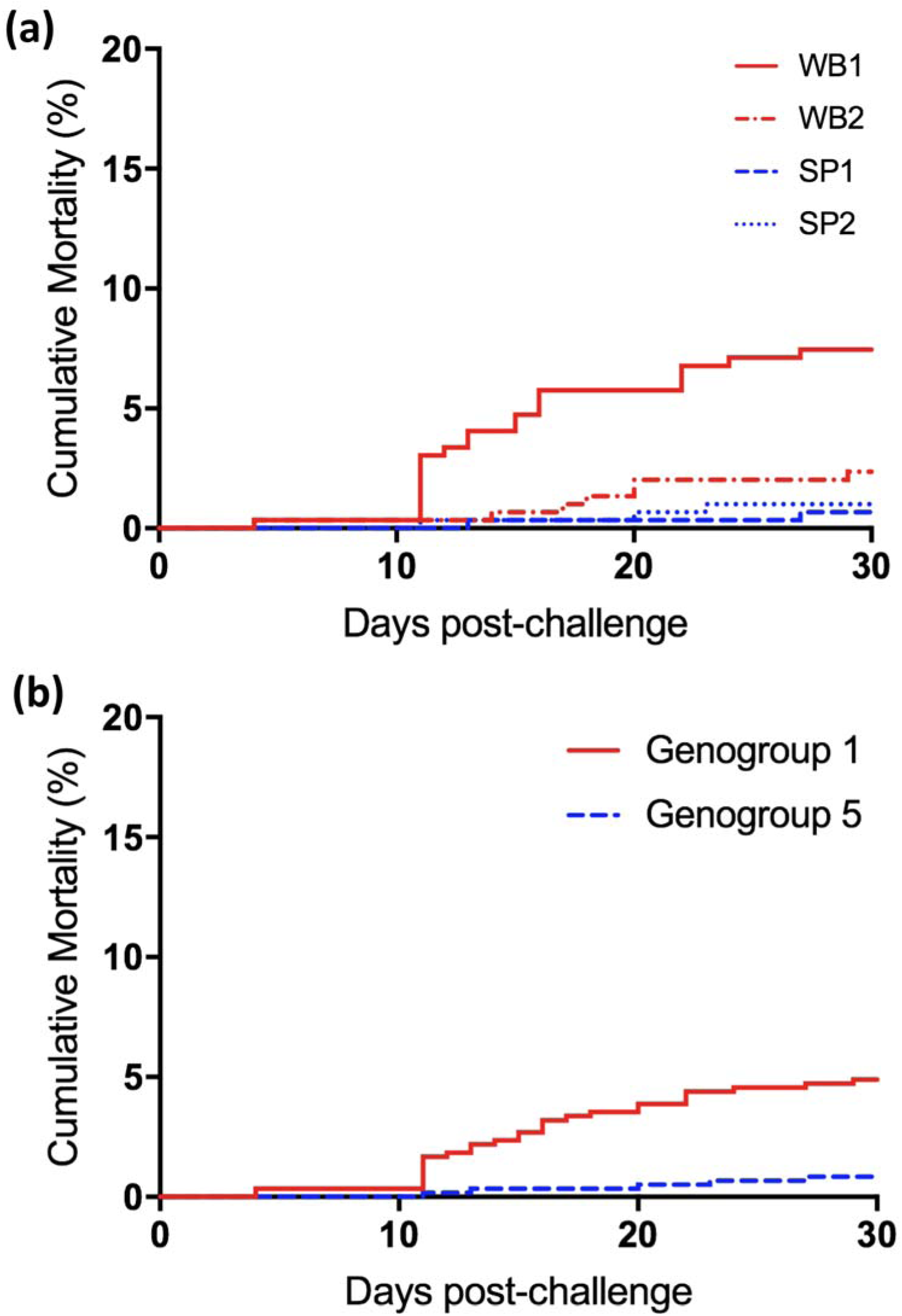
Kaplan-Meier mortality curves for rainbow trout infected with four isolates of IPNV from Genogroup 1 and 5. The cumulative mortality caused by each isolate is presented in a), and the combined mortality caused by isolates of each Genogroup is presented in b).

### IPNV testing and viral load

#### Mortality samples

A total of 49 mortality samples were tested to confirm the presence of IPNV by RT-qPCR and IFAT. The virus was detected in 34 of the analyzed samples (69.4%). IPNV positive samples by IFAT showed cells with a distinctive fluorescent staining, indicating that these samples contained infective viral particles. Samples ranged from a few fluorescent cells to hundreds per coverslip in high viral titer samples (Figure 2). There was substantial agreement between the two diagnostics methods (Kappa = 0.62, *p* < 0.0001), however, RT-qPCR was more sensitive, detecting IPNV in 33 of the samples tested, where only 26 positive samples were detected by IFAT. From the 34 positive samples, 29 were infected with isolates from genogroup 1, and 5 with isolates from genogroup 5. On average, virus load measured by RT-qPCR was 2 folds higher in fish infected with isolates from genogroup 5 than from genogroup 1. When the viral loads were compared by means of a ANCOVA using tank nested to genogroup as factor a statically significant difference was found (*p* = 0.005).

**Figure 2.**
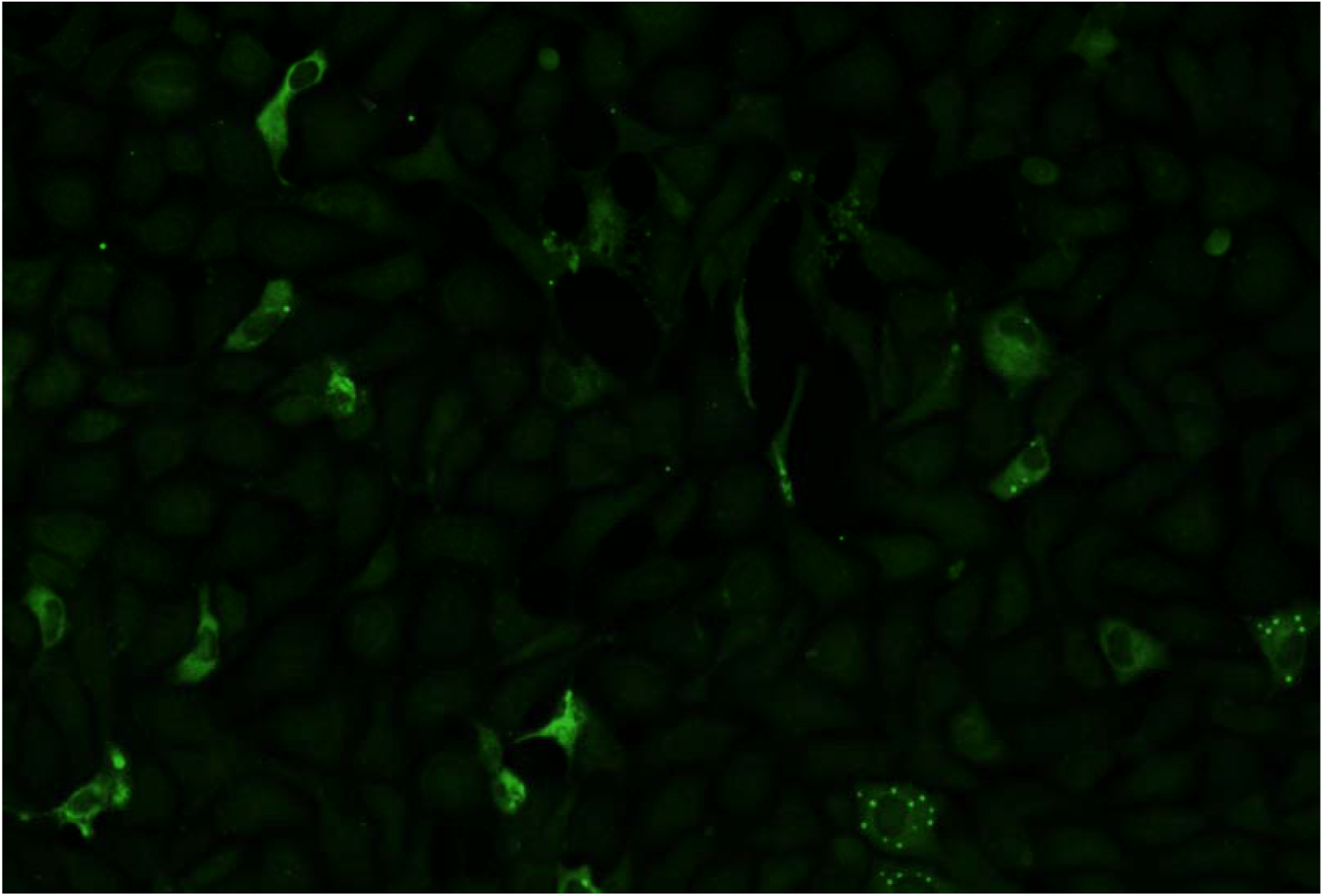
Indirect Fluorescent Antibody Test (IFAT) of IPNV positive mortality samples (10x). CHSE-214 cells infected with sample from fish challenged with Genogroup 5 isolate SP2 and stained with a polyclonal antibody against VP2 protein. Several infected (fluorescent) cells can be seen in the monolayer indicating a high viral titer.

### Survivor fish samples

Survivor fish sampled at day 1, 7 and 20 post infection, were analyzed to test for the presence of IPNV and to assess viral load by RT-qPCR. A total of 144 samples were tested and IPNV was found in 48 (33.3%), 23 belonging to fish infected with genogroup 1 isolates and 24 from genogroup 5. At day one post infection, IPNV positive samples were only detected in fish infected with isolate WB2, but showed very little VP1 mRNA expression, with a Ct value near cutoff value. Thus, one of these samples was set as calibrator for relative quantification of IPNV. At day 7 and 20 post infection all fish groups challenged with the different isolates showed at least one sample with the presence of the virus. Fish infected with isolate SP2 had the highest prevalence of IPNV, with 17 positive samples, while SP1 infected fish had the lowest, with seven samples positive to the virus (data not shown). In fish infected with isolates WB1 and WB2 from genogroup 1 the virus was detected in 12 and 11 of the analyzed samples, respectively. There was an increase in viral load throughout the challenge in fish infected with both genogroups, reaching a peak at day 20 post infection (Figure 3). As with mortality samples, viral load in survivor fish was, on average, higher in fish infected with isolates from genogroup 5 than with genogroup 1, and when the means of both groups were compared by an ANCOVA test, there was a statistically significant difference (*p* = 0.018).

**Figure 3.**
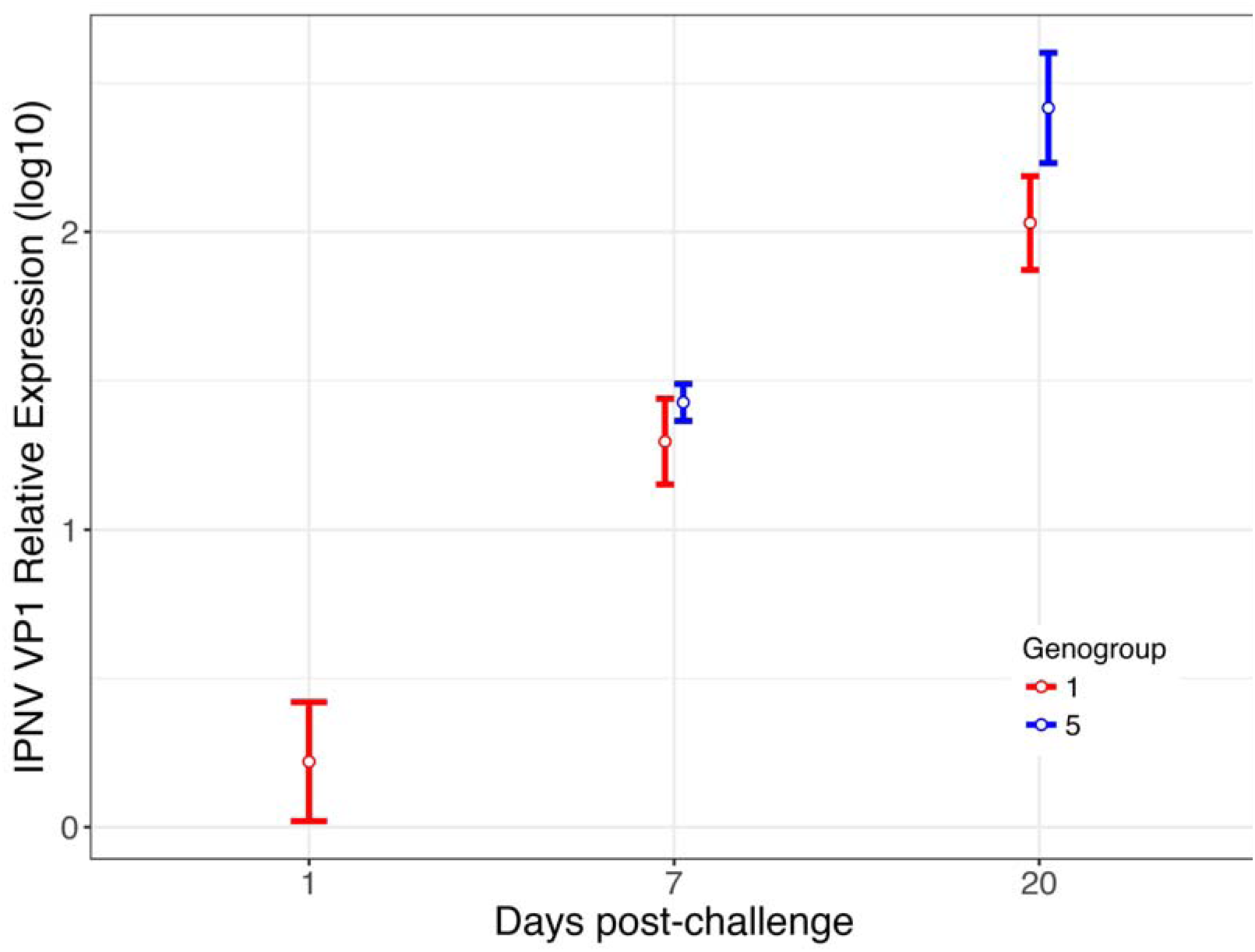
IPNV virus load in survivor fish sampled at days 1, 7 and 20 postinfection. Mean and standard error log10 VP1 transcript relative levels measured by RT-qPCR using ELF1α as housekeeping gene. No IPNV was detected in samples from genogroup 5 at day 1 post-infection.

## DISCUSSION

Previous reports have demonstrated that there is host genetic variation in resistance to IPN and that virulence may vary between IPNV isolates, causing great variation in mortality rates during outbreaks of the disease (Storset et al., 2007; Skjesol et al., 2011; Mutoloki et al., 2016). In this study, the mortality and viral load of rainbow trout fry individuals infected with Chilean IPNV isolates from genogroup 1 and 5 (more commonly known as strains WB and Sp, respectively) were compared by means of a 30-day experimental challenge. Our findings suggest that IPNV isolates from genogroup 1 were more virulent, i.e caused a higher mortality rate, than isolates from genogroup 5 in rainbow trout. This lends support to previous findings that suggested a host-specific relationship between IPNV isolates from genogroup 1 and salmonids from the genus *Oncorhynchus* farmed in Chile, based on the data from more than one hundred Chilean isolates from both genogroups (Tapia et al., 2015; Torres et al., 2016). Furthermore, since the publication of those studies these authors have continued to sequence more IPNV isolates, finding the same association between genogroups and salmonid species (Eissler et al. 2017). Conversely, Manriquez et al. (2017) recently suggested that there is no fixed relationship between salmonid species and genogroups of IPNV in Chile, since they found genogroup 1 isolates not only in *O. mykiss* and *O. kisutch* but also in *S. salar*. However, these authors molecularly characterized only 36 Chilean isolates, from which 10 where classified in genogroup 1, and only 4 came from *S. salar*. Hence, it is important to bear in mind that although there is a well known relationship between the genogroups of IPNV and salmonids present in Chile, this is not strict host specificity, and both salmon and trout can be infected with either genogroup of the virus.

This type of host-specific relationship has also been reported for other salmonid viruses, such as Infectious Hematopoietic Necrosis Virus (IHNV) in North America, where most of the isolates of the genogroups named U and M, come predominantly from sockeye salmon (*O. nerka*) and rainbow trout, respectively (Garver et al., 2003). Studies based on experimental challenges with both species and genogroups have shown that the relationship was associated with a specific virulence of IHN for each host, which depends mainly on the ability of the virus to enter the host fish and replicate (Garver et al., 2006; Peñaranda et al., 2009, 2011; Purcell et al., 2009). As with IPNV, IHNV isolates from both the U and M genogroups can infect rainbow trout; however, only the M genogroup viruses are highly virulent in this species. Using RT-qPCR they showed that viral load was significantly higher in trout infected with the more virulent M virus. Furthermore, a microarray analysis indicated that infection resulted in a greater overall host transcriptome change, suggesting that the M virus was more efficient at mediating host cell shutoff in order to enhance viral replication (Purcell et al., 2011).

Despite low mortality rates, viral load with IPNV genogroup 5 isolates was high and, these fish were indeed infected with IPNV and viral replication occurred until the last day of sampling, 20 days post infection. More interestingly, there is a tendency for genogroup 5 to have higher virus load on average than fish infected with isolates from genogroup 1, in both mortality and survivor fish samples. This would suggest that WB strain type isolates require the same or even less viral load than Sp train to cause greater mortality in rainbow trout. This is in contrast to what was seen in IHNV, and previous studies that show a positive relation between mortality caused by IPNV and virus load in Atlantic salmon, i.e. higher mortalities were associated with higher viral loads in susceptible fish or in fish infected with more virulent isolates (Skjesol et al., 2011; Reyes-Lopez et al., 2015; Robledo et al., 2016). It is important to point out that both isolates from genogroup 5 had contained high virulence marker, Thr_217_ and Ala221, in their VP2 sequence; whereas isolates from genogroup 1 had the avirulent motif, Pro_217_ and Thr_221_. Nonetheless, virulence was higher for genogroup 1 isolates in trout, suggesting that these virulence motifs are specific for the Sp strain, and that other genetic variants could have an effect on the virulence of the WB strain. Furthermore, in a previous challenge in Atlantic salmon, isolates SP1 and SP2 were able to cause significant mortality (data not shown), indicating that these viruses were virulent only in this species.

As was expected, mortality varied between fish infected with different isolates. However, overall mortality levels for the challenge were low, and only moderate differences in mortality were noted. It is well known that induction of overt IPN disease and mortality in experimental challenges is difficult to achieve, and is determined by several input variables like virus dose, fish age, temperature, infection route and host genetic susceptibility. For the challenge performed in this study we tried to meet the criteria for most of the variables recommended to obtain IPN-induced mortality in rainbow trout, by infecting first feeding fry via waterborne with a relatively high virus dose and maintaining them at 10 °C. However, it is plausible that a number of limitations could have influenced the results obtained and overall low mortalities reached in the challenge. One of the possible limitations was the virus dose.

Some authors recommend a dose as high as 1×10^7^ TCID_50_/mL per fish to attain higher mortalities in a cohabitation challenge with IPNV (Munang’andu et al., 2016). However, we were unable to obtain higher viral titers without increasing the number of passages of the isolates in cell culture. Nevertheless, several researchers have used the same dose used in this study in immersion challenges and have obtained considerably higher mortalities in rainbow trout and Atlantic salmon (Okamoto et al., 1987; Skjesol et al., 2011; Robledo et al., 2016).

Another technical limitation of this study is that neither phenotypic nor genotypic data for IPNV resistance of the rainbow trout individuals challenged was available. Recently it has been reported that mortality due to IPN in rainbow trout can range between 0% and 100% for the most resistant and susceptible families, respectively (Yoshida et al., 2019). Munang’andu et al. (2016) pointed out the importance of using highly susceptible fish to establish a challenge model for IPNV in Atlantic salmon that achieves mortalities above 75% in infected fish. Thus, it would be recommended that in future trials rainbow trout from known susceptible families are used in order to reach higher mortality levels that allow greater differentiation between groups.

Together these results show that IPNV isolates from genogroup 1 and genogroup 5 could infect rainbow trout but only the former caused important mortality in this species. This would suggest that IPNV isolates from the WB type strain are more virulent in rainbow trout than those from the Sp strain, and there is an association between the phylogenetic classification of IPNV and host susceptibility.

## ACKNOWLEDGEMENTS

The authors would like to thank Dr. Yoanna Eissler, Dr. Juan Kuznar and M.Cs. Juan Carlos Espinoza for their support in the completion of this study; the Laboratory of Virology of the University of Valparaíso for providing the viruses used and the infrastructure for the analysis of the samples; Germán Olivares and all the staff from the Rio Blanco hatchery for providing the experimental fish; and Veterquimica S.A. for enabling the facilities used in the challenge. We gratefully acknowledge the financial support provided by Subsecretaría de Pesca y Acuicultura (SUBPESCA) Proyecto FIP N° 2014-60 and Comisión Nacional de Investigación Científica y Tecnológica (CONICYT) scholarship 21161531.

